# Single TolC-AcrA complex formation monitored by time dependent single-channel electrophysiology

**DOI:** 10.1101/2022.06.14.496074

**Authors:** Igor V. Bodrenko, Tsedenia Alemu Zewdie, Jiajun Wang, Eshita Paul, Susanne Witt, Mathias Winterhalter

## Abstract

Characterizing protein-protein interaction on a single molecular level is a challenge, experimentally as well as the interpretation of the data. For example, Gram-negative bacteria contain protein complexes spanning the outer and inner cell wall being able to efflux effectively cell toxic substances. Within the search for antibiotics with novel modes of action, inhibition of the efflux pump assembly is an interesting possible target. Recent seminal work revealed the high-resolution structure of the tripartic composition TolC-AcrA-AcrB. Here, we reconstitute a single TolC homotrimer into a planar lipid membrane and follow the assembly of AcrA using the modulation of the ion current through TolC during binding. In particular, the presence of AcrA increases the average ionic current through TolC and, moreover, reduces the ion-current fluctuations caused by flickering of TolC. Here, we demonstrate that statistical properties of ion-current fluctuations (the power spectral density) provide a complementary measure of the interaction of the TolC-AcrA complex with potential inhibitors. Both characteristics, the average current and the current noise, taken into consideration together, provide more robust information.

## 1. Introduction

Characterization of the structure-function relationship of membrane proteins with their characteristic amphiphilic property requires a close to natural environment to avoid denaturation and precipitation. A particular class are channel forming proteins for which to obtain first structural information is to use cell patch clamp or alternatively, extraction, purification and reconstitution into artificial planar lipid membranes [1,2]. The modulation of the ion current across the membrane channel allows conclusion on channel gating, conformational changes or transport of small molecules [2]. The extreme contrast between the electric insulating membrane and a hydrophilic channel provides readily single channel resolution, however, the channel conductance as signal originates from the integral pathway of ions across the channel. Large channel conductance doesn’t imply large pore size. Even for large pore size the conductance is to a great extent controlled by the composition of the amino acids facing the channel lumen and the latter control substrate selectivity which can be also reversed by binding of divalent cations to carboxyl exposing amino acids (Aspartate, Glutamate) [2-5]. The amino acid composition may cause a huge internal electric field leading to orientation of the substrate’s dipole which in turn can favor or reduce permeation [3,5]. Examples for such observation are antibiotics permeating bacterial channels. Depending on the orientation of the dipole vs. the axis of the elongated antibiotic the permeation is drastically modified.

Most of the experimental set-up containing commercial amplifiers allow the recording of the ion current with sub millisecond resolution [2,6]. This range covers typical conformational changes but is not fully sufficient to detect small molecule permeation.

Bacteria are too small to be patched and for electrophysiological analysis the channels have to be extracted, purified and reconstituted into artificial planar membranes. Under favorable conditions also outer membrane vesicles containing overexpressed protein could be reconstituted [7]. Gram-negative bacteria contain a double cell wall separated by a peptidoglycan layer rendering them almost impermeable for antibiotics [8]. The outer membrane itself is rich in water-filled transmembrane proteins called porins that facilitate the passive selective permeation of small hydrophilic molecules. Interestingly, bacteria also may efflux toxic substances actively via the so-called efflux pumps among which is the *E. coli* TolC-AcrA-AcrB complex (for a recent special issue see [8]). Almost 20 years ago Koronakis and coworkers were able to provide a high-resolution structure of TolC revealing a hydrophobic membrane spanning beta-barrel porin combined with unusual long alpha helical loops reaching into the periplasmic space [9]. This large and long channel may bind to proton driven pumps located in the inner membrane. The connection between both protein units is supported by several AcrA bridging both into a functional unit. For all three subunits high-resolution structures were obtained and even more recently a high-resolution structure of the entire tripartite unit was obtained [10-13]. The TolC-AcrA-AcrB complex and his homologues have a multitude of functions and in particular are involved in antibiotic resistance as the complex can shuffle actively toxic compounds outside the bacteria. Most of the functional studies have been performed on a whole cell level.

Here, we focus on the assembly of AcrA to a single TolC channel. For this we insert a single TolC in a planar lipid bilayer and record the ion current across the channel. In the following step, we add AcrA while continuing to record the ion current. In a further series of experiments we test two well-known efflux pump inhibitors. The first one, hexamine cobalt binds to the tip of TolC and blocks the TolC channel effectively. A second inhibitor, PAβN, has several putative binding sites [14]. To test a possible inhibition of the complex formation we added PAβN before or after AcrA addition. In addition to the average ion current through the pore, we used the ion-current fluctuations (current noise) to get a complementary insight on the pore-substrate interaction.

## 2. Materials and Methods

### Expression and protein purification

TolC: For the protein expression, the pET24a based plasmid containing a C-terminally His-tagged full-length TolC was used as described previously [15] E.coli C43(DE3)ΔAcrAB cells, transformed with the TolC expression plasmid were cultured in TB media and in the presence of the appropriate selection marker at 37°C and 200 RPM to an OD600 of 1.3 and lysed using an Emulsiflex C3 Homogenizer (Avestin). After pelleting the cell debris, membranes were pelleted by centrifugation at 35000 RPM for 1h at 4°C. Membranes were resuspended and solubilized in loading buffer (20 mM Tris/HCl pH 7.5, 150mM NaCl, 10 mM Imidazole pH 7.5) at a concentration of 10 mL/g pellet in the presence of 1% DDM (Sigma). The solubilized membranes were cleared by centrifugation at 35000 RPM for 1h at 4°C. TolC was purified from the supernatant using a two-step chromatography approach on a BioRad FPLC system starting with an affinity chromatography step using an appropriate His-trap column according to the manufacturer’s manual followed by a size exclusion chromatography step using a S200 GL 10/300. All chromatography steps were performed in the presence of 0.03% DDM. Fractions containing trimeric TolC were pooled, concentrated and stored at 4°C until further use.

ArcA: The gene encoding ArcA without its signal peptide (amino acids 26 to 397) was amplified from an K12 derived E.coli strain and cloned into the NdeI and XhoI sites of pET21a. The integrity of the generated expression plasmid was confirmed by sequencing. For expression E.coli BL21(DE3)cells were transformed with the ArcA pET21a expression plasmid and grown in LB media and in the presence of the appropriate selection marker at 37°C and 180 RPM to an OD600 of 0.6 and subsequently induced with 1mM IPTG. Growth continued for 4 h at 25°C and 180 RPM. After cell harvest the pellets were flash frozen and stored at deep temperatures until further use. For cell lysis the cells were resuspended in 4 mL per g pellet in lysis buffer (20 mM Tris/HCl pH 7.5, 150 mM NaCl, 2 mM MgCl2, Protease Inhibitor (ROCHE), spatula DNAseI) and lysed using a sonotrode 106 Bandelin Sonicator. The soluble fraction was cleared by centrifugation for 30 min at 30.000xg and 4°C and applied to an appropriate His-trap column using an ÄKTA system according to the manufacturer’s manual. The affinity chromatography purification step was followed by size exclusion chromatography step using a S200 PG 16/600. Fractions containing dimeric ArcA were pooled, concentrated, aliquoted and flash frozen at deep temperatures until further use.

The Orbit Mini (Ionera, Freiburg) was used to form planar horizontal lipid bilayers [7]. A MECA 4 chip was first inserted, and a digital compensation was done after closing the lid. The acquisition frequency was set to 20 kHz. Afterwards, 150 µL of 1 M KCl and 10 mM HEPES electrolyte solution at pH 5.8 was added to the chip [15]. The bilayer can be formed on the four cavities using air bubbles. Firstly, a 10 µL micropipette was soaked in a lipid prepared from 1,2-diphytanoyl-sn-glycero-phosphatidylcholine at a concentration of 4-5 mg/ml in n-octane without aspirating any of it inside the pipette as there is enough residual lipid layer on the surface of the micropipette. The tip was placed near one of the cavities of the chip and a small air bubble was formed around it. The formation of the bilayer was assured by monitoring the current. After a bilayer was formed, around 0.1 mg/mL of TolC wild type was added to the chip. 100 mV was applied to assist the insertion of TolC into the bilayer. The insertion of a single channel corresponds to an increase of the current around 8-10 pA. The system was then diluted by a fresh buffer to avoid further insertions. An IV was measured for TolC and a gradient of concentrations (from 100 nM to 1250 µM) of several potential efflux pump inhibitor compounds were added and IV was measured accordingly. In another set of experiments, this time with AcrA, the TolC-AcrA complex was formed before adding the potential efflux inhibitor compounds. AcrA from a stock solution (3 mM) was added to obtain approximately 28 nM final concentration on top of the cavity that has the single TolC channel and incubated for 105 minutes.

The ion-current traces each between 40 to 150 second long were recorded in the ± 100 mV applied voltage range with the 25 mV step. The average current, its variance and the power spectral density (PSD) were calculated for each voltage value, using the algorithms described in [6].

## 3. Results

### 3.1 TolC single channel

In a first series of experiments we reconstituted a single TolC channel into horizontal planar lipid membranes as outlined in Materials and Methods. In order to have a reasonable resolution we used a 1 M KCl electrolyte solution and to obtain strong binding of AcrA to TolC we have chosen pH 5.8 [15]. Note that the set-up we used (Orbit Mini) allows only addition on the ground-electrode side and due to the strong shape asymmetry we assume that the α-helical part of the TolC faces the ground electrode.

In agreement with earlier measurements, our average current versus applied voltage gives (from the slope) an average conductance of 100 pS (1 M KCl, pH 5.8) for a single TolC channel (Fig 1. left)[16-19]. Note that using the Orbit Mini we often observed a few pA offset which is likely due to an electrodes offset or to some instrumental leakage in the cell. The IV curves are linear in the ±75 mV voltage range. In Fig. 1B, we show also the standard deviation of its noise at different applied voltage for four different experiments (Fig.1B). The current fluctuations show the voltage asymmetry, significantly increasing at +100 mV. We note that the absolute noise level might vary for each single-channel experiment, but it remains stable for different applied voltages.

**Figure.1.**
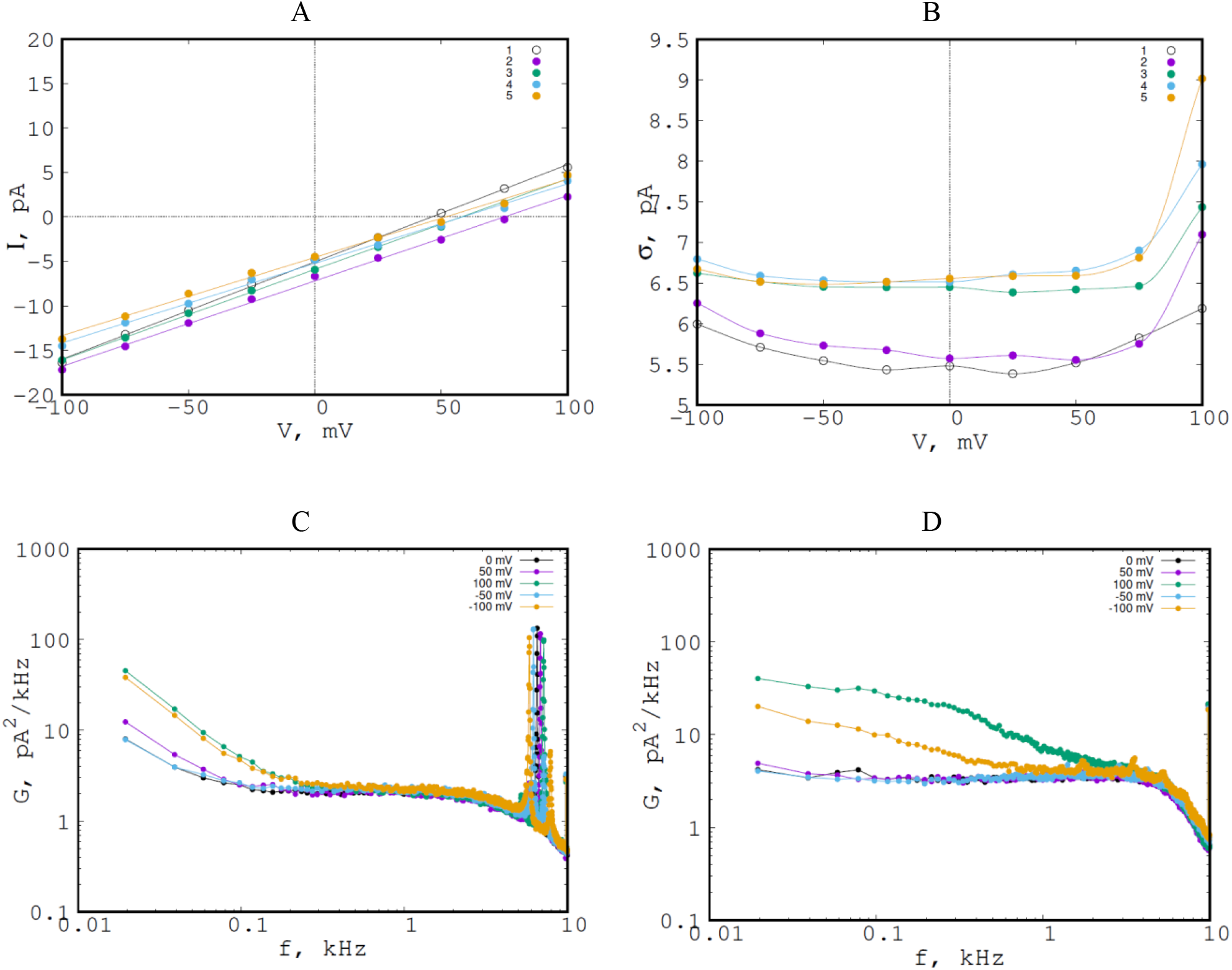
**A**. The average ion current (upper left) and **B**. the standard deviation of the current noise (upper right) versus applied voltage of a single TolC channel in 1 M KCl at pH 5.8. Each curve is the result of independent measurements. The points are the results of the measurements. The solid lines on the IV plot are the linear fits; the solid lines for the standard deviation are from the spline interpolation. **C, D** typical PSDs of the ion current through single TolC channels at different voltages for two independent reconstitutions, corresponding to the set 1 (left, C) and 2 (right, D), respectively, in fig 1A. The high-frequency peaks are due to induced external noise irrelevant to the studied process.

The PSDs of the ion current traces at different voltages obtained for two independent reconstitutions of the pore are shown on Fig. 1.C and 1.D, as typical examples. They demonstrate that both absolute value and a general shape of the PSD depend on the reconstitution (membrane and Tolc example). This implies that conclusions on the noise must be taken for each membrane preparation individually. The fluctuation data obtained for different reconstitutions can not be compared directly, as they characterize both the pore and the membrane. This is in contrast with the average current (the conductance), which is a characteristic of the pore only, as the membrane is an insulator.

The low-voltage conductance of TolC calculated for 24 independent single-channel reconstitutions of TolC are summarized in a histogram on Fig.2. It shows that TolC can probably be in different conformations stable during at least a few hours (the measurements time) and having the conductance in the range from 80 to 130 pS. The average conductance calculated over ol the data is 100 ps in agreement with previous experiments.

**Figure. 2.**
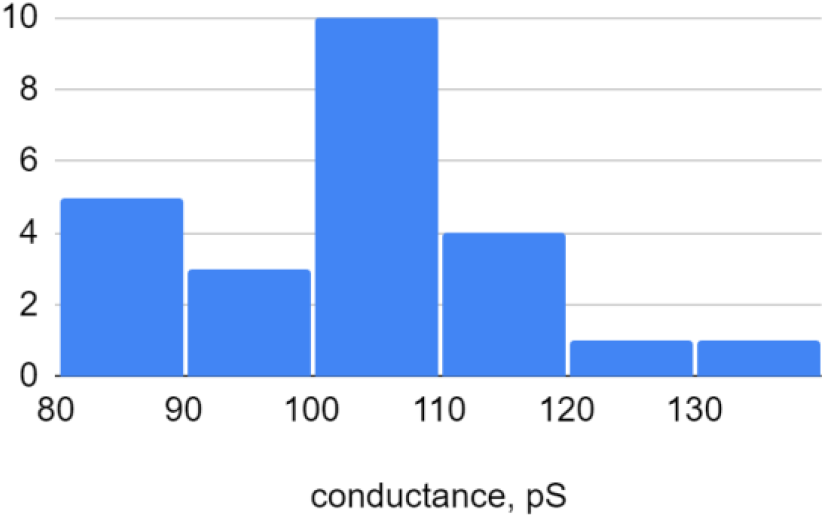
Histogram of occurrences of ion conductance values for 24 independent single-channel reconstitutions of TolC. The average current was 100 pS in 1 M KCl and at pH 5.8.

### 3.2 TolC-AcrA interaction

In the following series of experiments we study the effect of AcrA on the ionic current across TolC. For this, we benefit from the small volume of the Orbit Mini set-up which allows us to work with less than 40 µL of buffer. After reconstitution of a single TolC channel, we added AcrA to the α-helical/periplasmic side of the reconstituted TolC (see Materials and Methods).

The results of the analysis of the ion-current recordings made at various times after AcrA addition are shown on Fig 3. In **Fig.3A**, we notice that the conductance of the TolC+AcrA system increases during formation of the AcrA-TolC complex until it saturates after ca. 2h. We suggest that this time is required for diffusional redistribution of the added AcrA in the active volume of the cell. This is confirmed by 10 independent experiments with the TolC+AcrA complex.. On the other hand, the fluctuations of the ion current are systematically reduced after addition of AcrA (see **Fig.3B**). (Only in one case out of 10, the current fluctuations increased after addition of AcrC) Interestingly, this reduction of the fluctuations is due to the high-frequency (> 1 kHz) part of the PSD (Fig. 3C,3D).

**Figure. 3.**
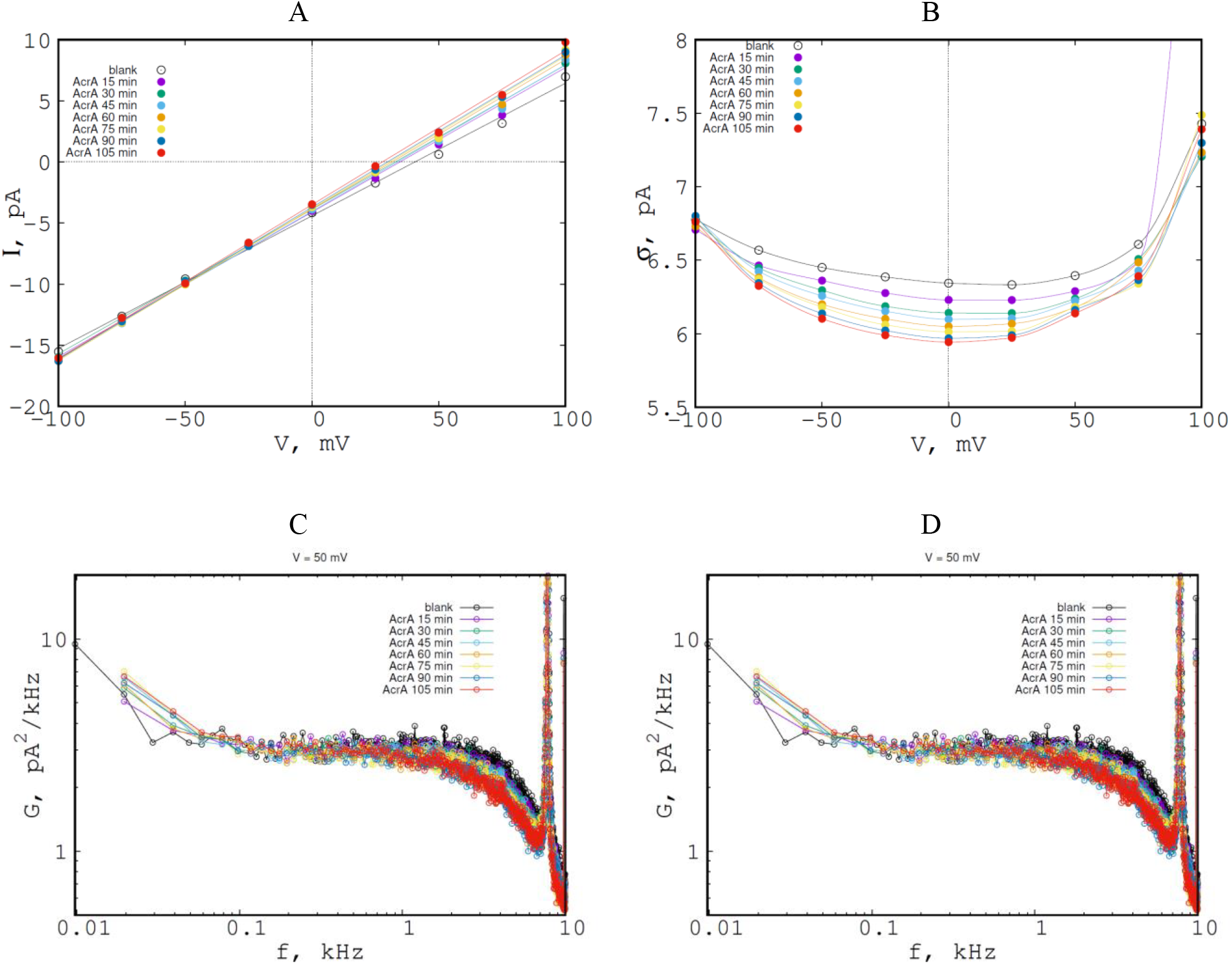
**A**. The average ion current (left) and **B**. the standard deviation of the current noise (right) through TolC single channel in presence of AcrA versus applied voltage; different curves are results recorded at different times after addition of AcrA; “blank” -only TolC. The points are the results of the measurements. The solid lines on the IV plot are the linear fits; the solid lines for the standard deviation are from the spline interpolation. **C**. Time dependence of the PSDs of the ion current through single TolC channels after addition of AcrA at +50mV (left) and **D**. -50 mV (right) of applied voltage. The data set labels correspond to those in A. The high-frequency peaks are due to induced external noise irrelevant to the studied process. Note the lowering of the noise in the 1-10 kHz range.

The low-voltage conductance of TolC and TolC+AcrA calculated for 10 independent single-channel reconstitutions of TolC are summarized in a histogram on **Fig.4**. One can suggest that AcrA stabilizes TolC in a higher conductance state. However, it does not increase the range of conductance, i.e. if the conductance of TolC was already high it was not increasing further after addition of AcrA (**Fig.4**.).

**Figure. 4.**
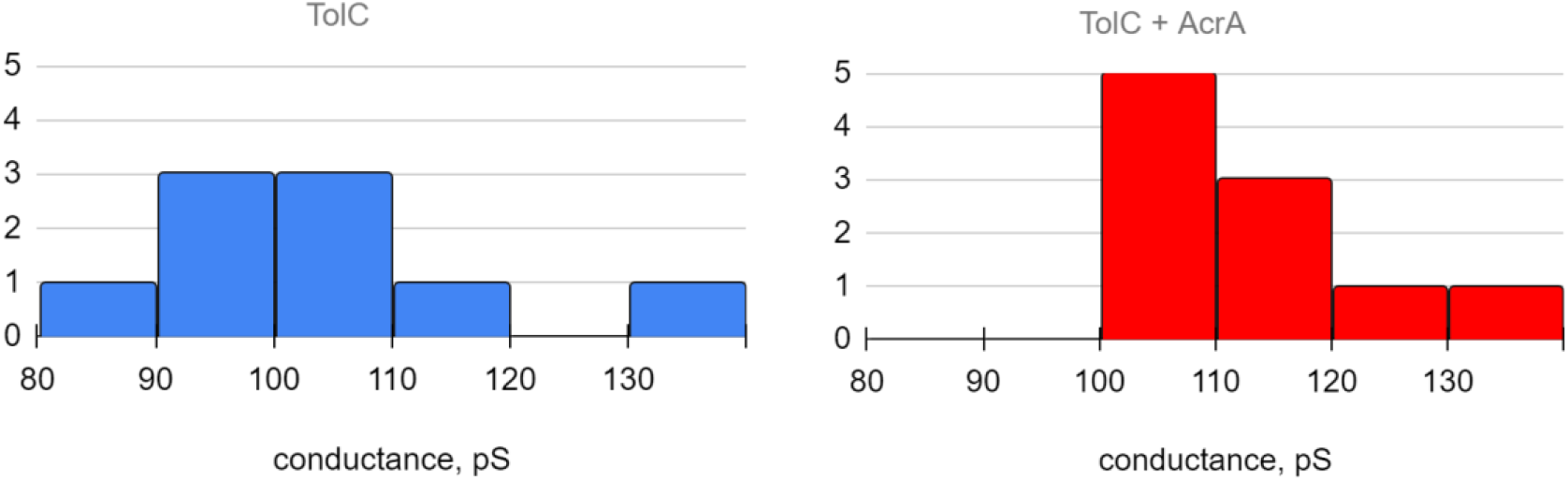
Ion current histograms for 10 independent single-channel reconstitutions of TolC (left) and after addition of AcrA (right). Note that addition of AcrA opens the channel (increases conductance). The buffer conditions were 1M KCl and pH 5.8.

### 3.3 TolC interaction with Hexamine cobalt (HC)

Several years ago Koronakis and coworkers showed that HC blocks the ion current through TolC blocks completely [17]. Inspection of the structure and molecular modeling suggest that HC binds via a strong charge-charge interaction at the tip of TolC and, as it sits in the middle of the channel, it is able to block the closed state. However to date, no significant biological inhibition was shown which might be simply caused by a low influx [17].

To examine the channel blocking ability, we titrate HC at the cis side of the chamber (corresponding to the periplasmic side of the TolC and also the side accessible for titration using the Orbit Mini). In Fig. 5 upper left, we show a typical I/V curve compared to a double mutant of wild type TolC replacing negative aa at the tip leading to a fully open channel. TolC alone is flickering with an open channel probability of about 70% under -100 mV bias voltage. Addition of 0.25 μM Hexamine cobalt decreased the open channel probability to 20%. By increasing the inhibitor concentration continuously to 1 μM, 5 μM, 10 μM, the open channel probability decreased to 8.2%, 5.3% and 1.2% respectively. These results are in agreement with the previous ones, and demonstrate the traditional effect of the HC binding in terms of the observable current-blockage effects.

**Figure. 5.**
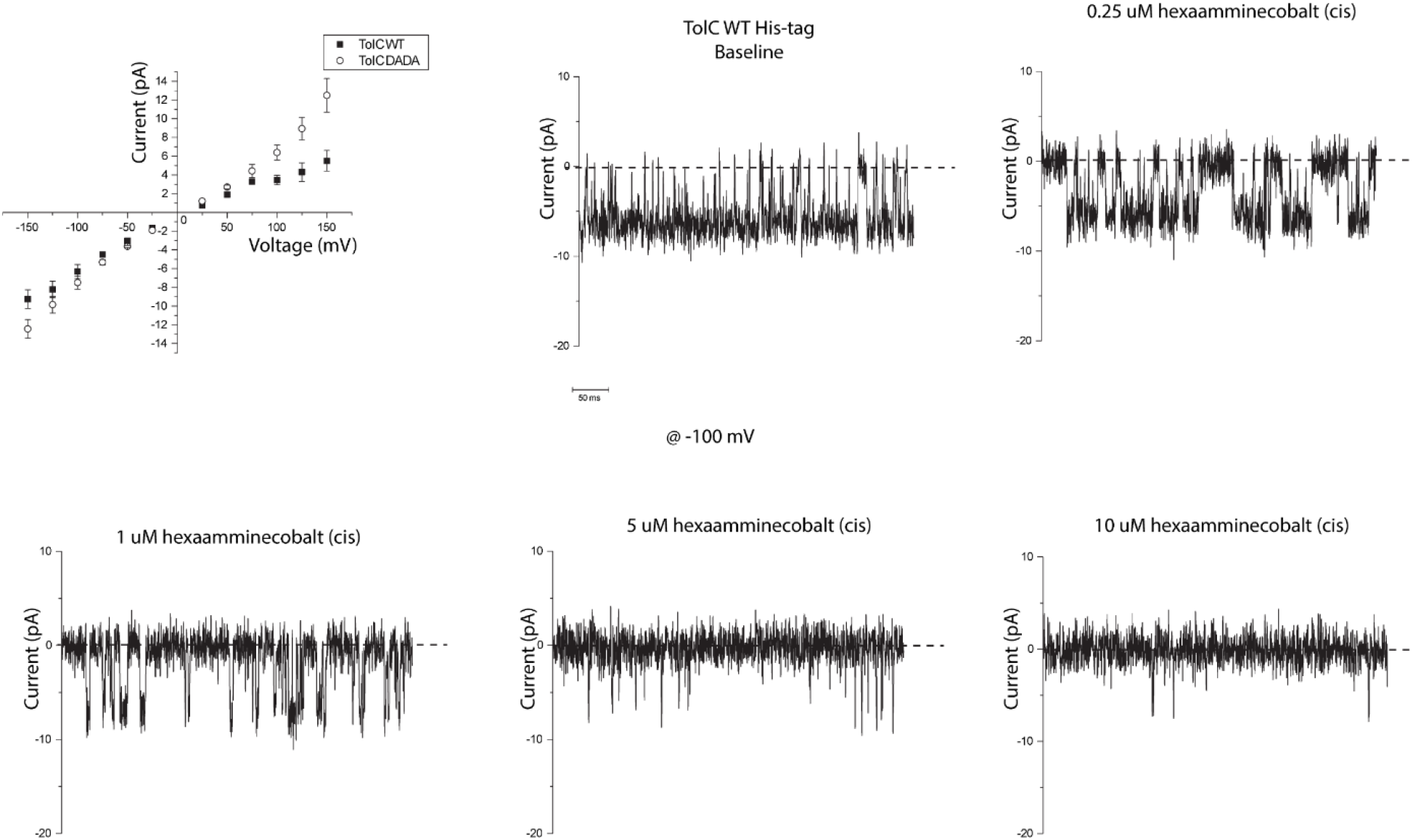
Interaction of hexamine cobalt (HC) with a single TolC channel. On the upper left the typical I/V curve for TolC wt and a mutant with removed negative aa at the tip, called DADA mutant is shown. For comparison we show the DADA mutant where the charge ring at the periplasmic exit has been replaced by alanine. The latter channel is mostly in the fully open form. The other figures show single channel traces with increasing HC concentration. The buffer contained 1M KCl at pH 5.8.

In an independent experiment, we study the time dependence of HC effect of the TolC, the HC was added (at 25 μM concentration) to the α-helical side of the reconstituted TolC. The results of the analysis of the recordings made at different times after the addition, immediately (few min), 15 min and 30 min, are shown in **Fig 6**. The stabilization of the results occurs after 15-20 min likely due to the low concentration of molecules requiring more time to reach the binding site. Thus, in the first recording the local concentration of HC at the mouth of the channel was smaller than less than 1 μM when the HC entrance into and the exit from the pore increases the current noise at low frequencies. After the concentration of HC in the cell is equilibrated at 25 μM (takes up to ca. 10-15 min) which corresponds to the saturation of the HC binding, the TolC channel is permanently blocked and the ion-current noise is reduced with respect to the one open channel.

**Figure. 6.**
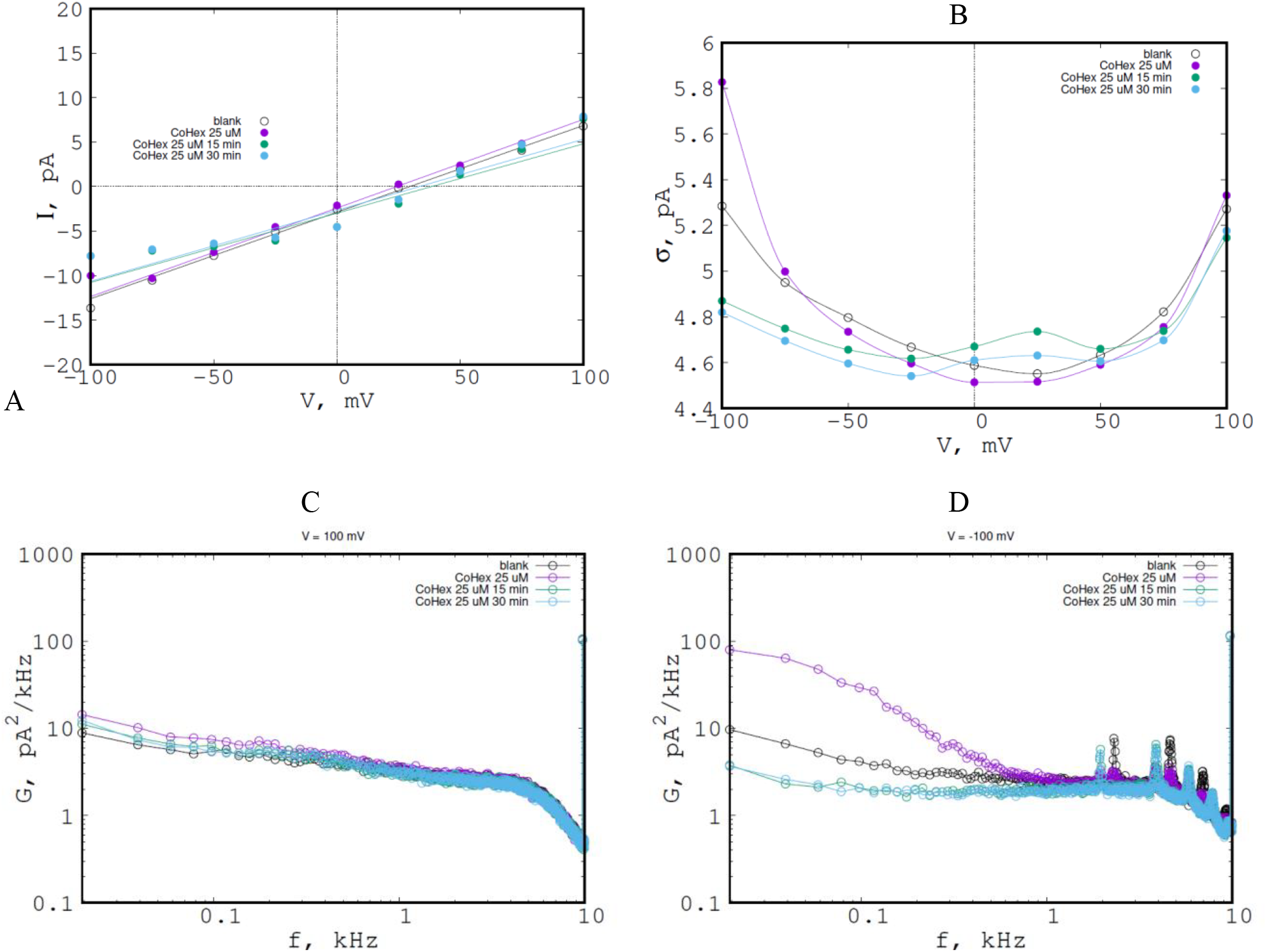
**A**.The average ion current (left) and **B**. the standard deviation of the current noise (right) through TolC single channel in presence of hexamine cobalt (HC) versus applied voltage; different curves are results recorded at different time after addition of CoHex; “blank” - only TolC.. The points are the results of the measurements. The solid lines on the IV plot are the linear fits; the solid lines for the standard deviation are from the spline interpolation. **6C**. Time dependence of the PSDs of the ion current through single TolC channels after addition of CoHex at +100mV (left) and **6D**. -100 mV (right) of applied voltage. The data set labels correspond to those in Fig.5. The high-frequency peaks are due to induced external noise irrelevant to the studied process.

At positive applied voltage, there is a very small effect of Hc on both the average current and the ion current noise. In contrast, at - 100 mV applied transmembrane voltage a dominant change at low frequencies (0.1-30 Hz) is observed corresponding to open-close transition. This transition is clearly different from the high frequency modulation visible in **Fig. 6**.

### 3.4 Effect of TolC inhibitor: L-Phenylalanin-β-naphthylamid (PAβN)

PAβN was reported in the literature to inhibit the function of the TolC-AcrAB efflux pump in the low μM range [8, 10, 14]. In order to elucidate a possible inhibition of the TolC-AcrA complex we investigated its effect on the ionic current through TolC. PAβN was titrated to the α-helical side of the reconstituted TolC and to the TolC+AcrA solution. The results are shown in **Fig.7**.

**Figure. 7.**
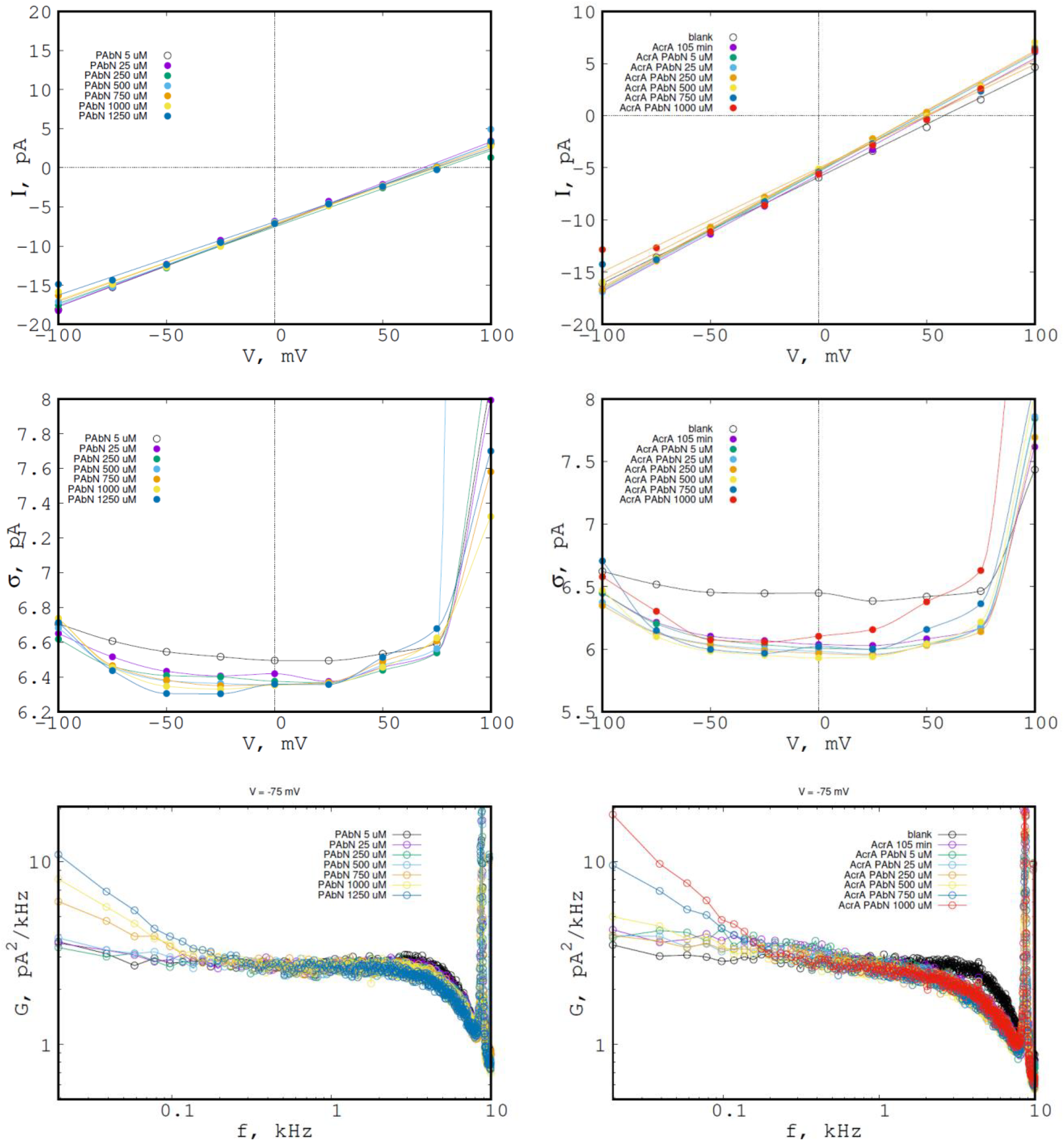
The ion-current characteristics in course of titration of PAβN to the TolC (left) and to the TolC+AcrA (right); “blank” - only TolC. The points are the results of the measurements. The solid lines on the IV plot are the linear fits; the solid lines for the standard deviation and the PSD plots are from the spline interpolation. The high-frequency peaks are due to induced external noise irrelevant to the studied process.

At low concentrations, 25-500 uM, PAβN reduces the ion-current noise of TolC due to reduction of the high-frequency (> 1 kHz) part of the PSD. Thus, it makes the effect similar to that of AcrA. In contrast, a low-concentration effect of PAβN on the TolC+AcrA complex is very small.

At higher concentrations, 750-1250 uM, there is a low-frequency (<100 Hz) increase of the PSD with concentration, both for Tolc and for TolC+AcrA. This behavior is in agreement with the hypothesis of the binding of PAβN to Tolc with at least partial blockage of the current, with the binding constant in the millimolar range. Interestingly, the events of current blockage are not well resolved on the current traces due to low conductance of the pore, high background noise and probably partial blockage.

## 4. Discussion

Most existing ion-current recordings have been done on planar lipid bilayer with accessible buffer volumes on both sides. Stable membranes can be obtained by various methods and typical volumes are several mL. For typical ion conductance recording this is feasible however titration with drugs such volumes are too large and miniaturization is needed. One commercial solution is the Orbit Mini, however, the active electrode is not accessible. This means, with respect to TolC, only the periplasmic side is available for titration. One advantage is the small volume, this allows one to use less than 150 μL or even as low as 40 μL of solution.

We reconstituted single TolC channels and characterized them by ion current recording. If TolC is added on one side only, this will lead to an asymmetry in the IV curve at higher voltage (> 100 mV) [16]. In agreement with previous results, we also observed a wide range, from 80 to 130 pS, of conductance values, suggesting different relatively stable conformations of TolC (of its flexible α-helical part). Besides, we have found that the voltage dependence of the ion current fluctuations is due to the low-frequency (< 1 kHz) processes, see Fig 1. The latter finding could be interpreted, for example, as the voltage driven transitions between the TolC conformations. These speculations, however, would require some evidence from computer modeling.

The low volume allowed titration of AcrA to TolC. As the effects of AcrA, we observed a small increase of conductance and a decrease of the ion.current noise, mostly due to the high frequency fluctuations (Fig.3, 4). We suggest that AcrA stabilizes the α-helical part of TolC upon binding, thus increasing the average ion current and reducing the current fluctuations.

In one control measurement, we titrated Hexamine cobalt to TolC. Being added to the periplasmic side, the HC demonstrates clear ion-current blockages in agreement with the previous measurements. The analysis of the current fluctuations (variance and PSD) demonstrate a characteristic increase of the fluctuations with the concentration of the substrate in the low-frequency (<100 Hz) part in agreement with a 2-state Markov binding process with the binding constant, K ca. 0.5 uM, and the correlation time in the 10 ms range.

In a further experiment, we studied the interaction of PAβN, a known efflux pump inhibitor, with TolC and TolC+AcrA. In this case, we could not clearly directly detect and quantify the ion-current blockages, due to the low pore conductance, high noise and probably incomplete current blockage. However, the current noise analysis could identify two effects of the PAβN. First, at 25-500 uM, PAβN reduces the ion-current noise of TolC due to reduction of the high-frequency (> 1 kHz) part of the PSD, similar to the effect of AcrA on TolC.

Besinds, at higher concentrations (> 750 uM) the noise increases at low frequencies, indicating partial channel blockage due to binding to TolC with the binding constant the millimolar range and the correlation time in the 10 ms range.

## 5. Conclusion

In the search for new antibacterial drugs, inhibitors of efflux pumps using TolC as a target are possible candidates. We have reconstituted a single TolC in a lipid bilayer and have studied its interaction with AcrA, (another efflux pump component), Hexamine cobalt (known TolC inhibitor), and PAβN (known efflux pump inhibitor) using single channel electrophysiology.

Addition of AcrA seems to open the channel slightly which is in agreement with the expectation from earlier work on the complex. A more careful inspection of the ion current traces revealed a strong flickering. It seems that binding of AcrA to TolC silences the tip of TolC, the putative binding site to AcrB to establish a fully functional efflux pump. We conclude that electrophysiology can contribute to mechanistic details of protein-protein assembly. The current noise analysis (variance and power spectral density) give complementary and often better insight into the pore-substrate interaction than the blocking-events counting in real time, especially in case of low conductance and high noise. A combination with fluorescence study is under way.

## 6. Acknowledgements

The research leading to these results was conducted as part of the Translocation consortium and has received support from the Innovative Medicines Initiative Joint Undertaking under Grant Agreement No. 115525, resources which are composed of financial contribution from European Union’s seventh framework program (FP7/2007–2013) and EFPIA companies. We further received support from the JPIAMR network RESET-ME (BMBF-ERANET JPIAMR - 01KI1827B) and the JPIAMR Virtual Institute Translocation-Transfer 01KI1828. Further support was provided by TSenArEO (BMBF-031B0864B). IVB acknowledges the financial support by DAAD (Deutscher Akademischer Austausch Dienst) during his research stay period at Jacobs University Bremen.

## 8. Highlights

- Binding of AcrA to single TolC channel modulates the ion current
- The TolC flickering is pronounced at 1-3 kHz.
- The periplasmic end of TolC is flexible, AcrA stabilize the open form
- Hexamine cobalt promoted the close state in a concentration dependent manner

## 9. TOC/Graphical abstract

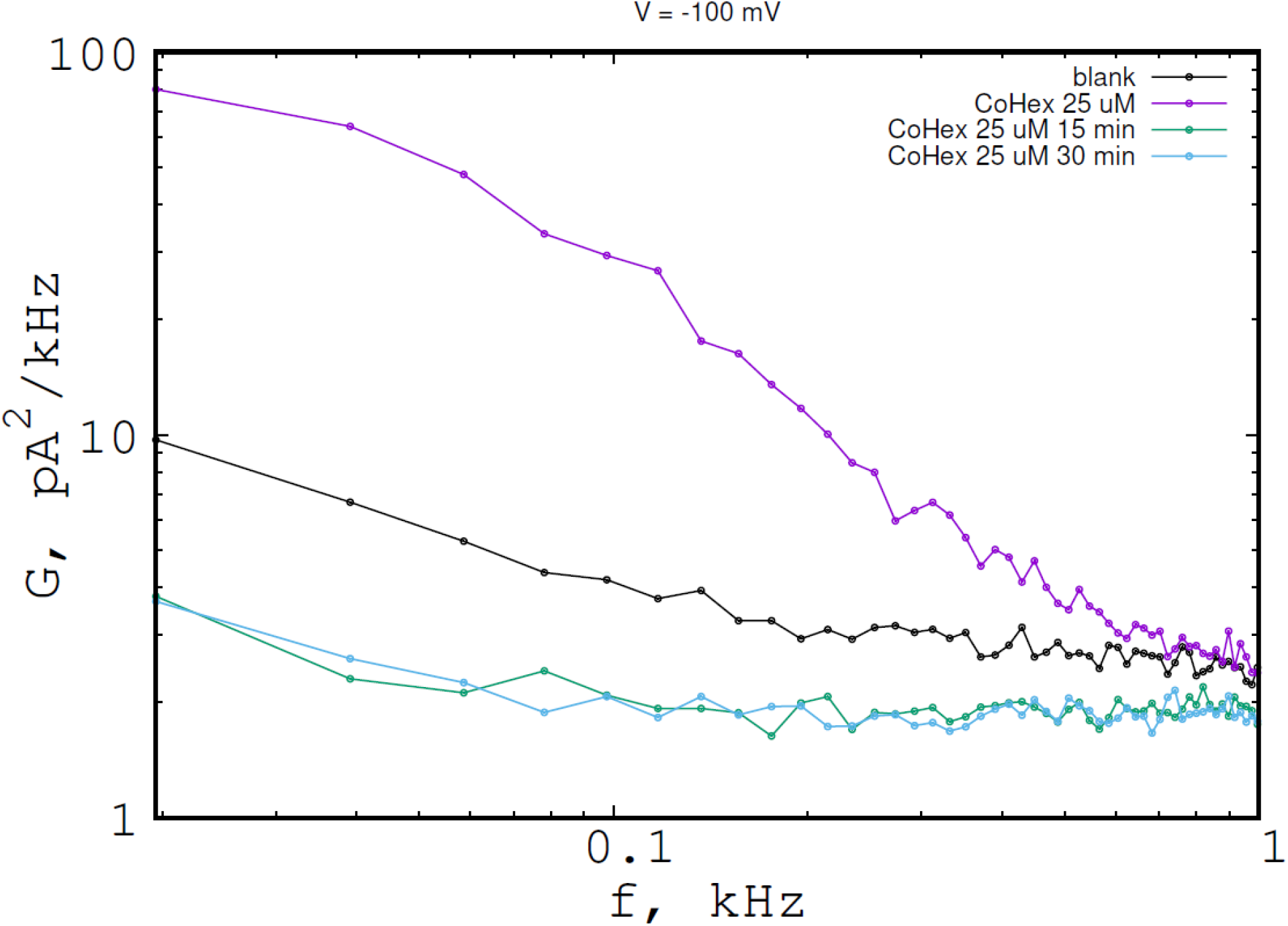

